# FRET-based Tau seeding assay does not represent prion-like templated assembly of Tau fibers

**DOI:** 10.1101/2020.03.25.998831

**Authors:** Senthilvelrajan Kaniyappan, Katharina Tepper, Jacek Biernat, Ram Reddy Chandupatla, Sabrina Hübschmann, Stephan Irsen, Sandra Bicher, Christoph Klatt, Eva-Maria Mandelkow, Eckhard Mandelkow

## Abstract

Tau aggregation into amyloid fibers based on the cross-beta structure is a hallmark of several Tauopathies, including Alzheimer Disease (AD). Trans-cellular propagation of Tau with pathological conformation has been suggested as a key disease mechanism. This is thought to cause the spreading of Tau pathology in AD by templated conversion of naive Tau in recipient cells into a pathological state, followed by assembly of pathological Tau fibers, similar to the mechanism proposed for prion pathogenesis. In cell cultures, the process is usually monitored by a FRET assay where the recipient cell expresses the Tau repeat domain (Tau^RD^, with pro-aggregant mutation, e.g., ΔK280 or P301L, ∼13.5 kDa) fused to GFP-based FRET pairs (YFP or CFP, ∼28 kD). Since the diameter of the reporter GFP (∼3 nm) is ∼6.5 times larger than the β-strand distance (0.47nm), this points to a potential steric clash. Hence, we investigated the influence of GFP tagged (N- or C-terminally) Tau^RD^ and Tau^FL^ (full-length Tau) on their aggregation behavior in vitro. Using biophysical methods (light scattering, atomic force microscopy (AFM), and scanning-transmission electron microscopy (STEM)), we found that the assembly of Tau^RDΔK^-GFP was severely inhibited, even in the presence of nucleation enhancers (heparin and/or pre-formed PHFs from Tau^RDΔK^). Some rare fiber-like particles had a very different subunit packing from proper PHFs, as judged by STEM. The mass per length (MPL) values of Tau^RDΔK^ fibrils are equivalent to 4.45 molecules/nm, close to the expected value for a paired-helical fiber with 2 protofilaments and cross-β structure. By contrast, the elongated particles formed by Tau^RDΔK^-GFP have MPL values around ∼2, less than half of the values expected for PHFs, indicating that the subunit packing is distinct. Thus, both kinetic and structural observations are incompatible with a model whereby external Tau can form a template for PHF assembly of Tau-GFP in recipient cells. As a consequence, the observed local increase of FRET in recipient cells must be caused by other signalling processes.

## Introduction

Tau, a microtubule-associated protein (MAPT, Uniprot P10636), has an important role in microtubule assembly and stabilization. Tau has a hydrophilic, mostly basic composition, is natively unfolded and is highly soluble. Nevertheless, Tau amyloidogenic aggregates characterize a wide range of neurodegenerative diseases known as Tauopathies (Wang and Mandelkow, 2016, Lee et al., 1988, Holtzman et al., 2016) including Alzheimer Disease (AD). Mutations in the Tau gene alone are sufficient to cause neurodegeneration (Hutton et al., 1998). Moreover Tau deposits in the brain correlate well with the memory decline, confirming the importance of Tau pathology in AD (Braak stages) (Nelson et al., 2012, Braak and Braak, 1991). Biophysical and structural studies show that soluble monomeric Tau, upon nucleation by polyanionic cofactors like heparin or RNA, can form insoluble paired helical filaments (PHFs) in vitro (Kampers et al., 1996, Goedert et al., 1996). However, the pathways causing Tau aggregation in neurons and of Tau-induced neurodegeneration are not well understood.

In AD, Tau pathology spreads from the entorhinal cortex to anatomically connected regions such as hippocampus, subiculum and cortex. The spatio-temporal progression of cognitive impairment correlates well with the Tau pathology, as assessed by hallmarks such as aggregation or hyperphosphorylation (Braak and Braak, 1991, Braak et al., 2006). This has led to the hypothesis that the disease progression in AD is caused by the cell-to-cell spreading of Tau protein itself in a pathological state (Holmes et al., 2014, Kfoury et al., 2012). In support of this hypothesis, Tau can be secreted from neurons (concentration in ISF ∼1 nM (Yamada et al., 2011)), secretion is enhanced by neuronal activation (Pooler et al., 2013, Wu et al., 2016), by exosomes (Wang et al., 2017) and by neuronal death (Arriagada et al., 1992). Extracellular Tau can be taken up by neighboring cells by several mechanisms including receptor mediated endocytosis, phagocytosis, muscarinic receptor-mediated or HSPG mediated uptake (Michel et al., 2014, Gomez-Ramos et al., 2008, Holmes et al., 2013). The internalized Tau is thought to induce the fibrous aggregation of endogenous Tau by templated self-assembly. This would promote further aggregation and propagation Tau pathology to other cells, by analogy to the mechanism proposed for prion pathogenesis (Colby and Prusiner, 2011). Based on the assumption that spreading of Tau protein is responsible for the spreading of neuronal pathology, current therapeutic approaches include the prevention of the pathological conformation of Tau, scavenging extracellular Tau by antibodies, blocking of Tau uptake by neurons, reducing Tau concentrations, and others (DeVos et al., 2017, Wobst et al., 2015, Mirbaha et al., 2015, Yanamandra et al., 2013).

The key methods for investigating the reactions of cellular Tau protein in response to external Tau are based on expressing aggregation-prone forms of Tau repeat domains (Tau^RD^) labeled with fluorescent sensors, e.g. CFP, YFP. Their accumulation can be observed by local increases of fluorescence, FRET, or FLIM which indicate proximity within several nm (Furman and Diamond, 2017, Kfoury et al., 2012, Vaquer-Alicea and Diamond, 2019, Gorlovoy et al., 2009). The repeat domain (RD) of Tau represents the assembly-competent core of Tau filaments, as it contains the hexapeptide motifs with a high propensity to generate β-structure (von Bergen et al., 2000, Wischik et al., 1988). Mutations such as ΔK280, P301L, P301S in the repeat domain enhance β-propensity of Tau and hence show stronger toxic effects (Mocanu et al., 2008, Santacruz et al., 2005). An increased FRET signal in a recipient cell is usually taken as a sign of pathological PHF-like Tau aggregates, induced by the pathological conformation of the Tau seeds penetrating into the cells. This would amount to a prion-like propagation of Tau pathology from cell to cell (Holmes et al., 2014, Kfoury et al., 2012, Frost et al., 2009).

From a structural perspective, the templated assembly of Tau into bona fide PHFs can only be concluded if the Tau aggregates are filaments with cross-β structure (axial repeat between strands ∼0.47nm), similar to PHFs from AD brain (Fitzpatrick et al., 2017). We hypothesized that a large GFP reporter molecule (∼28 kD) tagged onto Tau^RD^ (∼13 kD) could inhibit the aggregation because of a steric clash: The diameter of the reporter GFP (∼3 nm) (Yang et al., 1996) is ∼6.5 times larger than the β-strand distance (0.47nm) between the Tau molecules. To test this, we investigated the assembly forms of Tau^RD^ tagged with GFP either at their C-terminus or N-terminus, using several biophysical and microscopic methods. For comparison we also studied the assembly forms of GFP-tagged full-length Tau, and the same Tau proteins without GFP tags. The results showed that the self-assembly of Tau^RD^ is severely inhibited when tagged with GFP. In particular, even when fiber-like particles occurred they had a very different structure and mass distribution, distinct from that of PHF-like Tau fibers. We conclude that cell inclusions with enhanced FRET intensity do not result from a templated assembly of PHF-like Tau filaments, and do not result from the transfer of a pathological conformation.

## Results

### Generation of Tau constructs to study the effect of fusion protein (GFP) on Tau

For in vitro studies, GFP-tags were fused to either full length Tau protein (2N4R) or the shorter repeat domain containing the “pro-aggregant” deletion mutation of K280 (Tau^RDΔK^) at their N-terminus or C-terminus using a flexible linker (Fig. 2A, B). The proteins were expressed in *E. coli*, except Tau^RDΔK^-GFP which could only be expressed in the baculovirus-Sf9 cell system. His-tags were added at the C-termini to aid purification.

**Figure 1:**
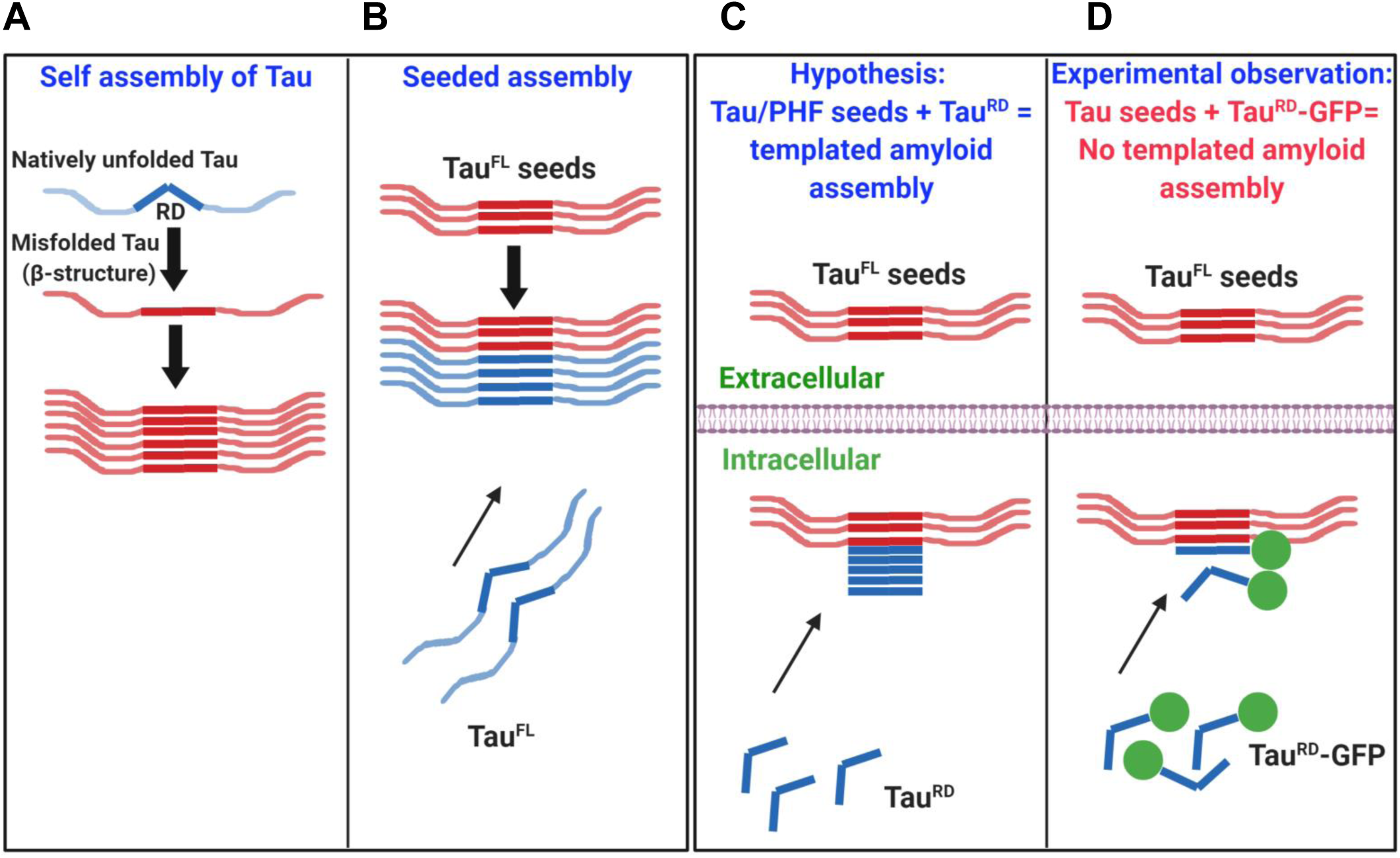
Schematic representation of effects of GFP label in preventing templated assembly of Tau ^RD^. **(A)** Self-assembly of Tau: Tau protein is highly hydrophilic and natively unfolded. In disease conditions, Tau becomes misfolded, attains β-structure and assembles into paired helical filaments. **(B)** Seeded (templated) assembly: Tau protein subunits elongate into long filaments using external Tau seeds as template (usually prepared by sonication of preformed fibrils). Note: In Tau assembly experiments in vitro, polyanions such as heparin must be added to initiate nucleation. **(C)** Pre-dominant hypothesis: In cell culture, Tau seeds (made from Tau^FL^ or Tau^RD^) are internalized into acceptor cells. Endogenous Tau^RD^ is thought to attain the amyloid structure of the seeds and elongate the filaments. **(D)** Experimental observation: In cell culture studies Tau^RD^-GFP is often used as endogenous Tau fusion protein, intended to elongate templating seed structures. However, steric hindrance by the large GFP molecule prevents fiber formation.

**Figure 2:**
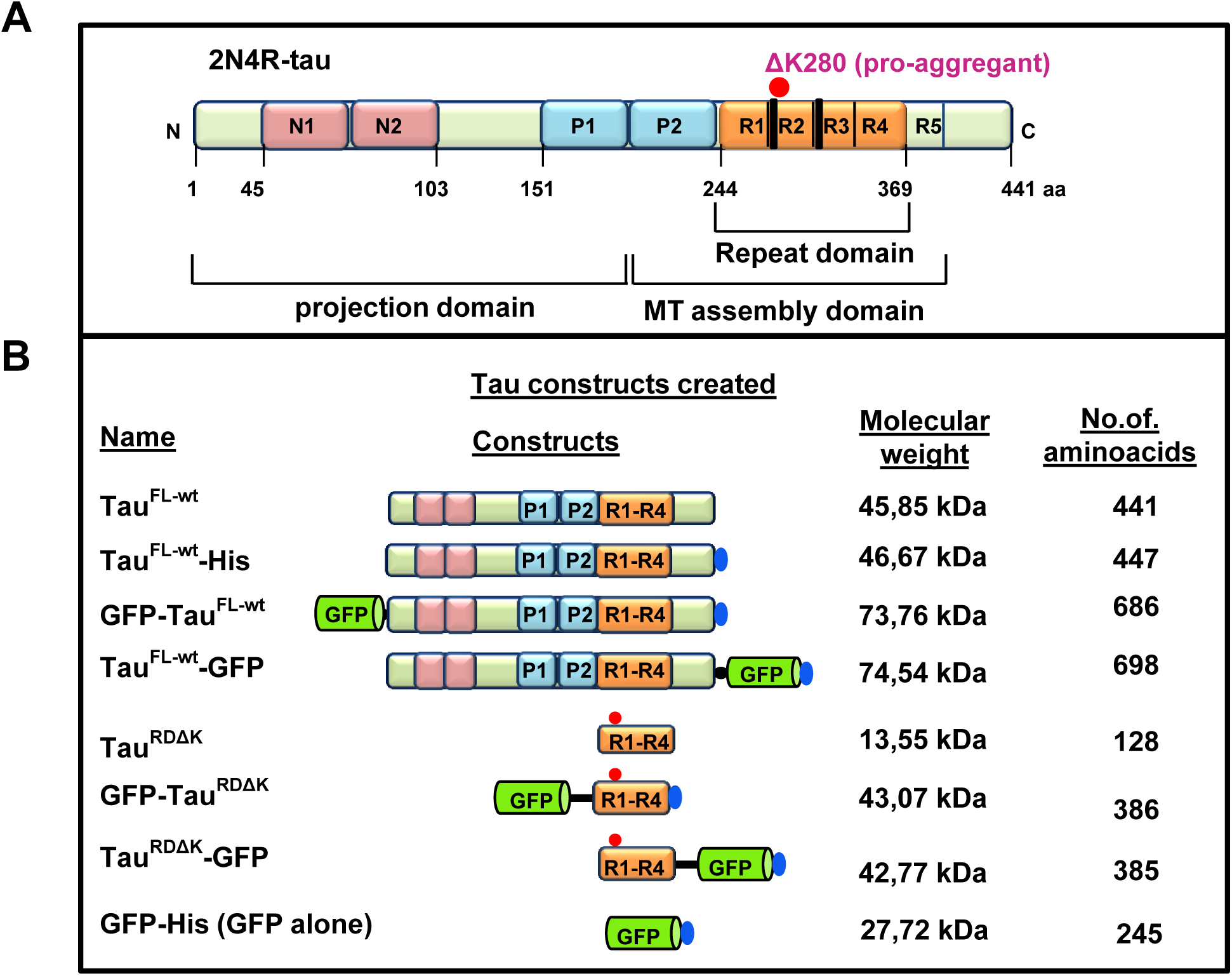
Tau proteins and variants used in this study. **(A)** Bar diagram of longest isoform of Tau in human CNS comprising 441 amino acids (Uniprot P10636-F, alias htau40, Tau-2N4R). Domains are depicted with 2 near-N-terminal inserts (N1, N2, pink), the proline-rich domains (P1, P2; blue), 4 pseudo-repeats (R1-R4, ∼31 residues each, ochre, plus one less-conserved repeat R5), and the C-terminal tail. The amyloidogenic hexapeptide motifs are indicated at the beginning of R2, R3 (thick black line). The C-terminal half (MT assembly domain P2-R5) promotes both microtubule assembly and pathological aggregation of Tau, the N-terminal half (projection domain) projects from the surface of microtubules or from the core of Tau fibers, respectively. The “pro-aggregant” deletion mutation ΔK280 (red dot) lies near the beginning of R2, it increases the β-propensity of Tau and hence increases pathological aggregation. **(B)** Tau and GFP fusion proteins are schematically presented with their name listed on the left. The GFP-tag is positioned either at the N-terminus or C-terminus. Some constructs contain a C-terminal 6xHis-tag to aid in the purification (blue ellipse). A sequence of 13-14 amino acids was used as a linker (black line) in the two short Tau constructs GFP-Tau^RDΔK^ and Tau^RDΔK^-GFP to increase the distance between the GFP and the repeat domain (see Table S1).

### GFP does not interfere with microtubule assembly induced by full-length Tau

We first asked whether GFP fusion to Tau affects the physiological function of promoting microtubule assembly as monitored by light scattering at 350 nm (Gaskin et al., 1975) and verified by electron and fluorescence microscopy (Fig. S1A-C). Full length Tau, either untagged or labeled with GFP at either end, was competent to induce MT assembly with similar time courses (t_1/2_ ∼ 1-2min, Fig. 3A, top curves). GFP alone remains soluble, Fig. 3A, bottom curve). By contrast, the Tau repeat domain alone, either with or without ΔK mutation, and with or without GFP tag, is not competent to cause MT polymerization (Fig. 3B, bottom curves). This is consistent with the weak binding of Tau^RD^ to tubulin and microtubules, which becomes pronounced only when the domains flanking the repeats are present (Gustke et al., 1994). The results mean that GFP moieties attached to the either end of full-length Tau are sufficiently flexible and/or far from the MT binding domain to allow assembly without steric hindrance.

**Figure 3:**
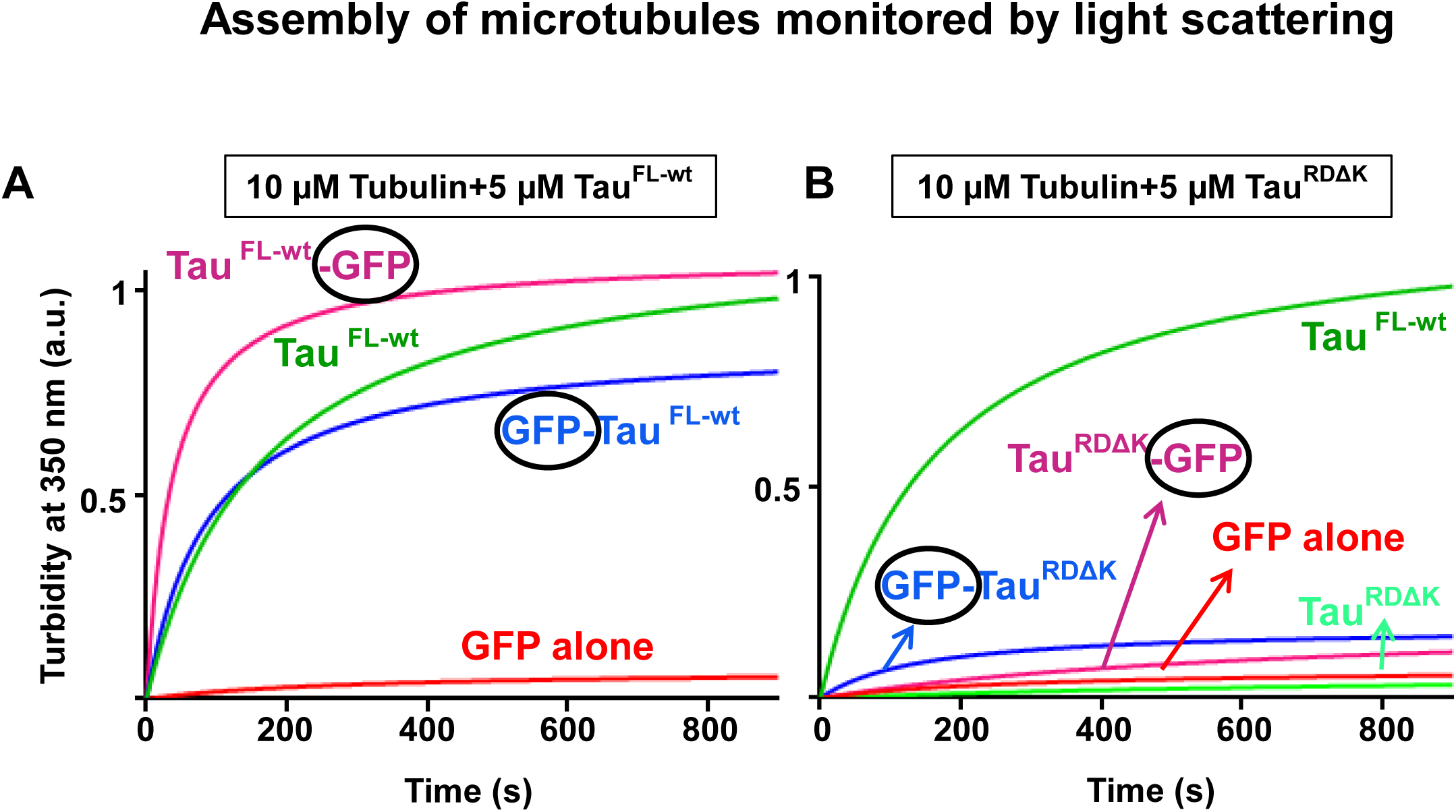
Full length wildtype Tau proteins (with or without GFP) induce microtubule assembly whereas the repeat domain does not. **(A)** Assembly of microtubules (10µM tubulin) is supported by Tau^FL-wt^ (5µM) without GFP or with GFP attached to either N- or C-terminus (top, green, blue, magenta curves). MT assembly levels and rates (t_1/2_ ∼min range) are comparable. Note that GFP alone shows no increase in light scattering (red curve). **(B)** Microtubule assembly is not supported by the repeat domain Tau^RDΔK^, regardless of GFP (bottom curves: Tau^RDΔK^ (light green); GFP-Tau^RDΔK^ (blue); Tau^RDΔK^–GFP (magenta); GFP alone (red)). As a control, full length Tau^FL-wt^ induces robust microtubule assembly (as in Fig. 3A, green). Note that the repeat domain R1-R4 is often denoted as “MTBD” domain in the literature, even though it binds and assembles microtubules poorly.

### GFP tags create steric hindrance against aggregation of Tau repeat domain

The central question of this paper is to see whether GFP fusion of the Tau repeat domain allows aggregation into bona fide PHF-like filaments. To test this, Tau^RDΔK^ (Fig. 4B) and Tau^FL-wt^ Fig. 4A) with and without GFP were incubated with the nucleating reagent heparin16000 at 37°C and aggregation was monitored by UV light scattering at 350 nm. In the case of the repeat domain, the overall extent of aggregation of Tau^RDΔK^ (Fig. 4B, green) was 3-fold higher than that of Tau^RDΔK^-GFP (magenta) whereas GFP-Tau^RDΔK^ (blue) showed almost no response, indicating that GFP attached to either end of the Tau repeat domain was strongly inhibitory, even though it carried the pro-aggregant mutation ΔK280. A different picture emerges from full length Tau: Here the unlabeled and GFP-labeled proteins showed a robust increase of light scattering during assembly (Fig. 4A), with some variation in assembly rates and final levels. Note that GFP increases the Tau subunit mass by ∼28kD/48kD∼58%, roughly consistent with the increase in scattering.

**Figure 4:**
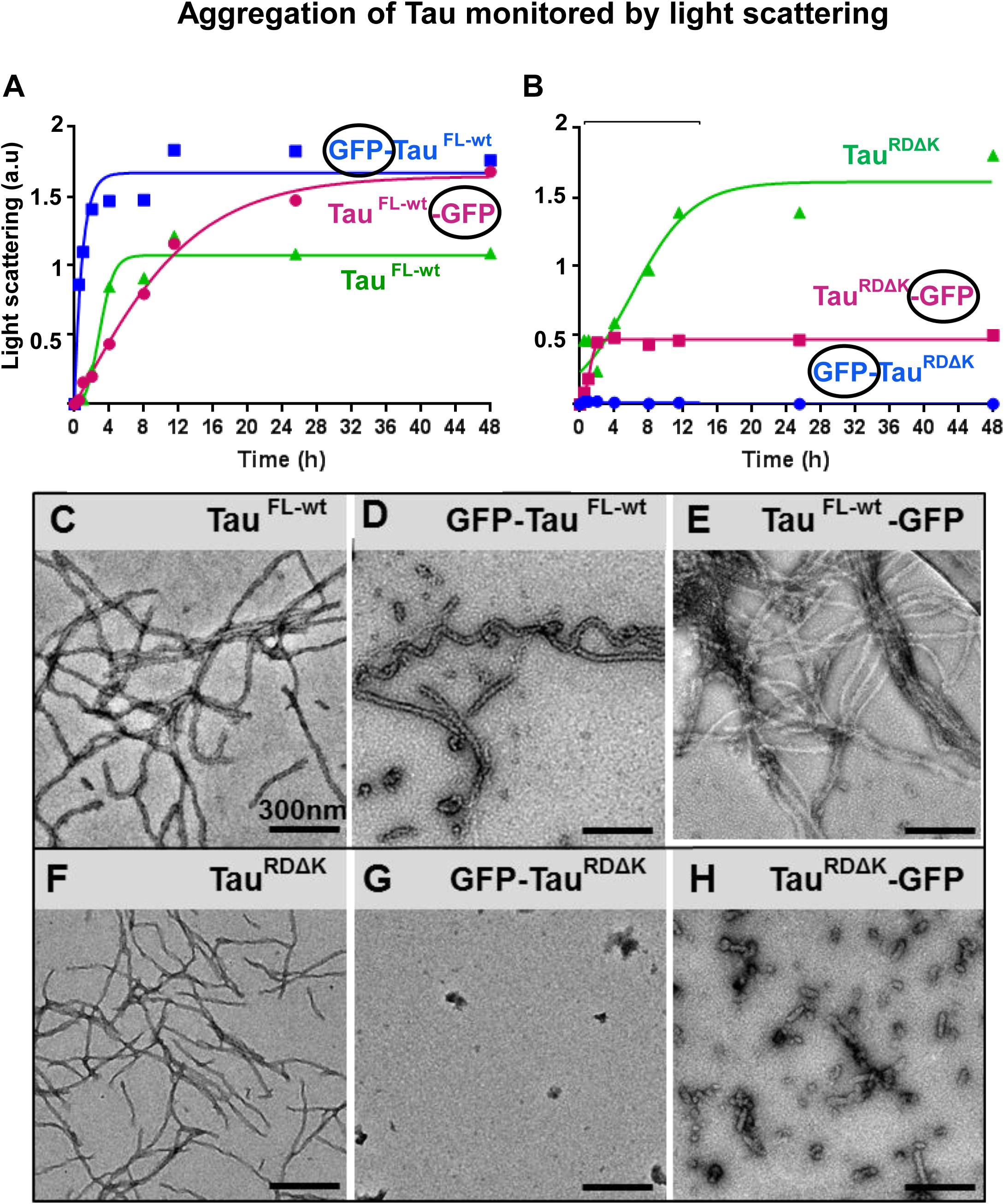
Aggregation of Tau proteins with GFP tags monitored by light scattering and electron microscopy. **(A-B) Attachment of GFP to Tau repeat domain severely inhibits aggregation, but full length Tau+GFP remains aggregation competent**. Aggregation of Tau to PHF-like fibers was monitored by light scattering at 350 nm. 50µM of full length Tau^FL-wt^ or Tau^RDΔK^ with or without GFP on either end was incubated in the presence of heparin 16000 (12.5µM) in BES buffer, pH 7.0, at 37°C at different time intervals. **(A)** Unlabeled Tau^FL-wt^ (green) aggregates mostly into PHF-like fibers with t_1/2_ ∼ 3 h. GFP–Tau^FL-wt^ (blue, t_1/2_ ∼2 h) and Tau^FL-wt^–GFP (magenta, t_1/2_ ∼12 h) aggregate with distinct rates and reach higher final levels, consistent with the higher mass of the subunits and variations in elongation. **(B)** Unlabeled Tau^RDΔK^ (green curve) shows aggregation with t_1/2_ ∼6 h into PHF-like fibers. By contrast, GFP-Tau^RDΔK^ (blue) shows no increase in light scattering, and Tau^RDΔK^–GFP (magenta) reaches only a low level of light scattering saturating at ∼2h. Thus in both cases the GFP-labeled protein is severely inhibited in aggregation. **(C-H) Electron microscopy reveals heterogeneous aggregation forms of Tau+GFP** Samples of Tau^FL-wt^, Tau^RDΔK^ with and without GFP aggregated for 24 h at 37°C were placed on carbon grids and imaged by negative stain electron microscopy. Top row, full length Tau^FL-wt^ proteins with or without GFP forms typical long filaments (C-E), whereas repeat domain Tau^RDΔK^ does not (F-H, bottom row). **(C)** Tau^FL-wt^-GFP aggregates into typical PHF-like twisted filaments (diameter ∼28 nm). **(D)** GFP-Tau^FL-wt^ forms long filaments with increased diameter (∼40 nm). **(E)** Tau^FL-wt^-GFP forms long filaments and bundles (∼28 nm) and filament bundles. **(F)** Repeat domain Tau^RDΔK^ without GFP forms filaments of typical PHF-like morphology but smaller diameter (∼21 nm). **(G)** GFP-Tau^RDΔK^ does not form filaments but only amorphous small aggregates. **(H)** Tau^RDΔK^-GFP forms short filaments and oligomers with the length <100nm; average diameter ∼37 nm.

### GFP fused Tau^RDΔK^ forms amorphous small aggregates, oligomers and aberrant filaments

The data from light scattering and sedimentation assays (data not shown) were correlated with structural investigations using electron and atomic force microscopy. Full length Tau^FL-wt^ aggregated into long (straight and twisted) fibrils (apparent diameter ∼24 nm, Fig. 4C). GFP-Tau^FL-wt^ also formed straight or twisted filaments (Fig. 4D) but appeared somewhat thicker in negative stain (∼40 nm), consistent with the extra mass of GFP. Tau^FL-wt^-GFP formed long, straight or twisted fibrils with a tendency to coalesce into bundles (Fig. 4E). As a control, the His-tag on Tau^FL-wt^-^His^ had no influence on assembly (similar to Fig 4C, data not shown).

More pertinent to the present study are the results on the shorter construct Tau^RDΔK^ which readily forms PHF-like fibers with β-structure (Fig. 4F, see Barghorn et al., 2000). However, in contrast to full-length Tau, filament formation strongly inhibited by GFP-tags on Tau^RDΔK^. GFP fused to the N-terminus (GFP-Tau^RDΔK^) completely abolished fiber aggregation, and only small amorphous assemblies were generated (Fig. 4G). GFP fusion to Tau^RDΔK^ at C-terminus (Tau^RDΔK^-GFP) made it difficult to purify in E.coli due to protein instability, and hence the protein was expressed in Sf9 cells. This protein formed predominantly oligomers and few elongated particles with thicker diameters, distinct from PHFs (∼37 nm, Fig. 4H).

### Atomic force microscopy reveals GFP decoration around the core of fibrils assembled from full-length Tau-GFP

As an alternative imaging approach we used AFM in tapping mode of unstained Tau aggregates. The spatial resolution in the x-y-plane is limited by the size of the tip, but the z-height is recorded accurately via the tip touching the protein surface. Fig. 5 shows overview images of Tau filaments (left), magnified views (middle), and plots of the height distribution across filaments. Tau^FL-wt^ shows long twisted or straight filaments (Fig. 5A) with heights of ∼8 nm and apparent widths of ∼42 nm, enlarged ∼2-fold by the tip-broadening effect, compared with the actual width 10-25 nm (Wegmann et al., 2010). Filaments of GFP-Tau^FL-wt^ (Fig. 5B) show a similar core (height ∼7nm), but in addition a surrounding halo of height ∼3nm which matches the size of the GFP moiety.

**Figure 5:**
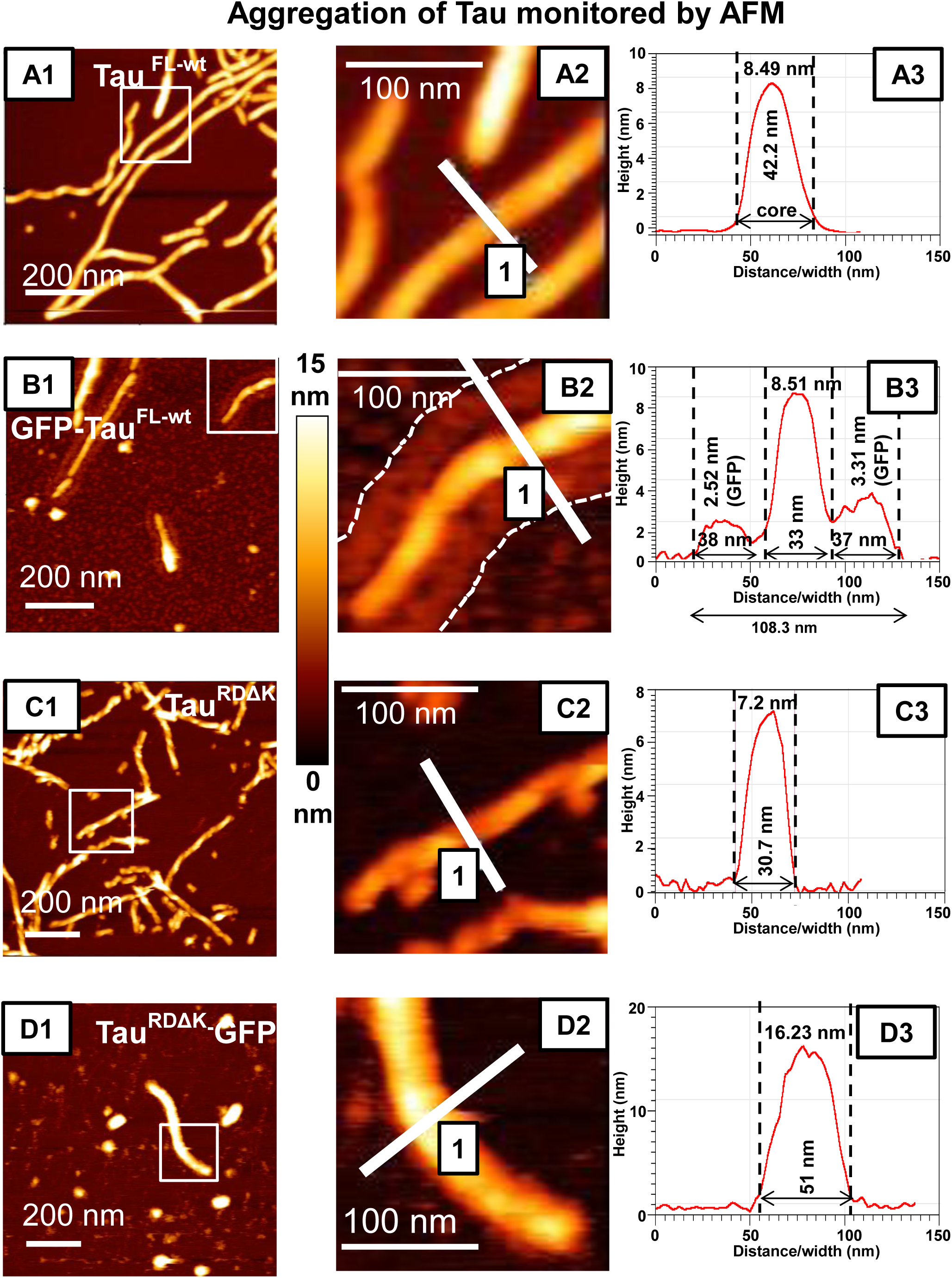
Atomic force microscopy reveals the changes in Tau filament assembly and decoration with GFP. AFM analysis of Tau fibrils formed in the presence of heparin for 24 h at 37°C was imaged in tapping mode using an MSNL cantilever. Left row, overviews, center row, magnified details, right row, line scans showing distribution of height. **(A1, B1)** Tau^FL-wt^ **(A1)** and GFP-Tau^FL-wt^ **(B1)** both form long fibrils (straight or twisted), but the height distribution shows characteristic differences because the attachment of GFP generates a halo on both sides of the filament core (compare A3 and B3, core and side bands indicated by dashed lines. The average height (thickness) of the fibril core is similar for Tau^FL-wt^ (7.7 nm± 0.7 nm; n=36) and GFP-Tau^FL-wt^ (7.0 nm± 1.2 nm; n=36). However, the overall thickness increases from Tau^FL-wt^ alone (38.1 nm± 5.45 nm; n=50) to GFP-Tau^FL-wt^ (96.01 nm± 8.98 nm; n=50). The two the side peaks (B3) are ∼3nm wide, in good agreement with the shape of GFP. (**C-D**) The repeat domains construct Tau^RDΔK^ forms well twisted filaments **(C1)** whereas Tau^RDΔK^-GFP forms only few short filaments and more globular shaped aggregates **(D1)**. The enlarged images **(C2, D2)** show that the short filaments of Tau^RDΔK^-GFP are thicker than those of Tau^RDΔK^. (**C3** and **D3)** The widths and heights of Tau^RDΔK^-GFP (width - 42.95 nm± 8.32 nm; n=50; height-18.5 nm± 3.1 nm; n=36) fibrils are larger than those of Tau^RDΔK^ fibrils (width - 28.17 nm± 3.35 nm; n=50; height - 7.0 nm± 1.1 nm; n=36). The height scale for A1, B1, C1 and D1 is 0 to 15nm.

Unlike GFP tagged full length Tau, Tau^RDΔK^-GFP aggregates are mostly small oligomers and some short thick filaments with height of ∼16nm (Fig. 5D), about twice the value of unlabeled fibers (Fig. 5C).

Taken together, the kinetic and structural data show that attachment of GFP to Tau^RDΔK^ on either side severely inhibits its aggregation, and the small fraction of fiber-like aggregates is distinct from PHFs.

### Mass-per-length analysis by STEM discriminates PHF-like filaments from aberrant structures

PHFs from Alzheimer brains consist of two protofilaments with cross-β structure (Fitzpatrick et al., 2017). Given the axial spacing between β-strands of ∼0.47 nm, the molecular density of protein subunits would be 1/0.47=2.13 units per nm in each protofilaments, or ∼4.26 molecules per nm in a PHF, irrespective of the molecular weight of the subunits, and thus the mass-per-length (MPL) should be ∼4.2 times the subunit MW. The MPL value can be determined by STEM which allows one to distinguish PHF-like subunit packing from aberrant structures. Tau filaments with and without GFP were prepared at 37°C for 24 h for STEM analysis and the MPL data were compared with standard MPL data of tobacco mosaic virus (TMV). The results (Fig. 6 and Table 1) show that fibers assembled from the unlabeled repeat domain Tau^RDΔK^ agree very well with the theoretical expectations (∼4.4 molecules/nm). Fibers from unlabeled full-length Tau^FL-wt^ have ∼20% lower values, ∼3.4 molecules/nm, even though their appearance by EM or AFM is similar to those of the repeat domain. Both observations are in excellent agreement with our earlier study using a different set of instruments and protein preparations (von Bergen et al., 2006). The apparent decrease by ∼20% can be explained by the fact that the “fuzzy coat” of full-length Tau is spread out around the perimeter so that its contrast is partly buried in the background. On the other hand, in the case of GFP-labeled proteins, the small fraction of fiber-like structures had very different subunit packings, e.g. ∼0.9 or 2.0 for the “long” and “short” fibrils of Tau^RDΔK^-GFP, and ∼2.2 for GFP-Tau^FL-wt^ fibrils. These values are clearly incompatible with a PHF-like packing of molecules, yet their GFP moieties are close enough to generate FRET signals. MPL data evidently show that GFP fusion changes the aggregation pattern and packing of Tau molecules in filaments (Table 1 for detailed MPL values).

**Table 1.** Summary of the effects of Tau + GFP fusion proteins on aggregation, stimulation of microtubule assembly, and packing of subunits in Tau filaments

| Construct | Expression<br>(Cell type) | MT<br>assembly | Aggregation<br>(+Heparin) | Mol. weight<br>(kDa) | MPL<br>(kDa/nm) | No.of. molecules<br>/nm |
| --- | --- | --- | --- | --- | --- | --- |
| Tau<sup>FL-wt</sup> | *E.coli* | yes | Fibrils | 45.85 | 157.1±4 | 3.4 |
| GFP-Tau<sup>FL-wt</sup> | *E.coli* | yes | Fibrils | 73.76 | 163.0±5 | 2.2 |
| Tau<sup>FL-wt</sup>-GFP | *E.coli* | yes | Fibrils | 74.54 | n.d. |  |
| Tau<sup>RDΔK</sup> | *E.coli* | no | Fibrils | 13.55 | 61.5±4 | 4.5 |
| GFP-Tau<sup>RDΔK</sup> | *E.coli* | no | no | 43.07 | n.a. |  |
| Tau<sup>RDΔK</sup>-GFP | Sf9 cells | No | Short fibrils<br>(minor fraction) | 42.77 | 86.1±5 | 2.0 |
|  |  |  | Long fibrils<br>(minor fraction) | 42.77 | 37.4±7 | 0.87 |
| TMV |  |  |  |  | 131.1±1 kDa (standard) |  |

**Figure 6:**
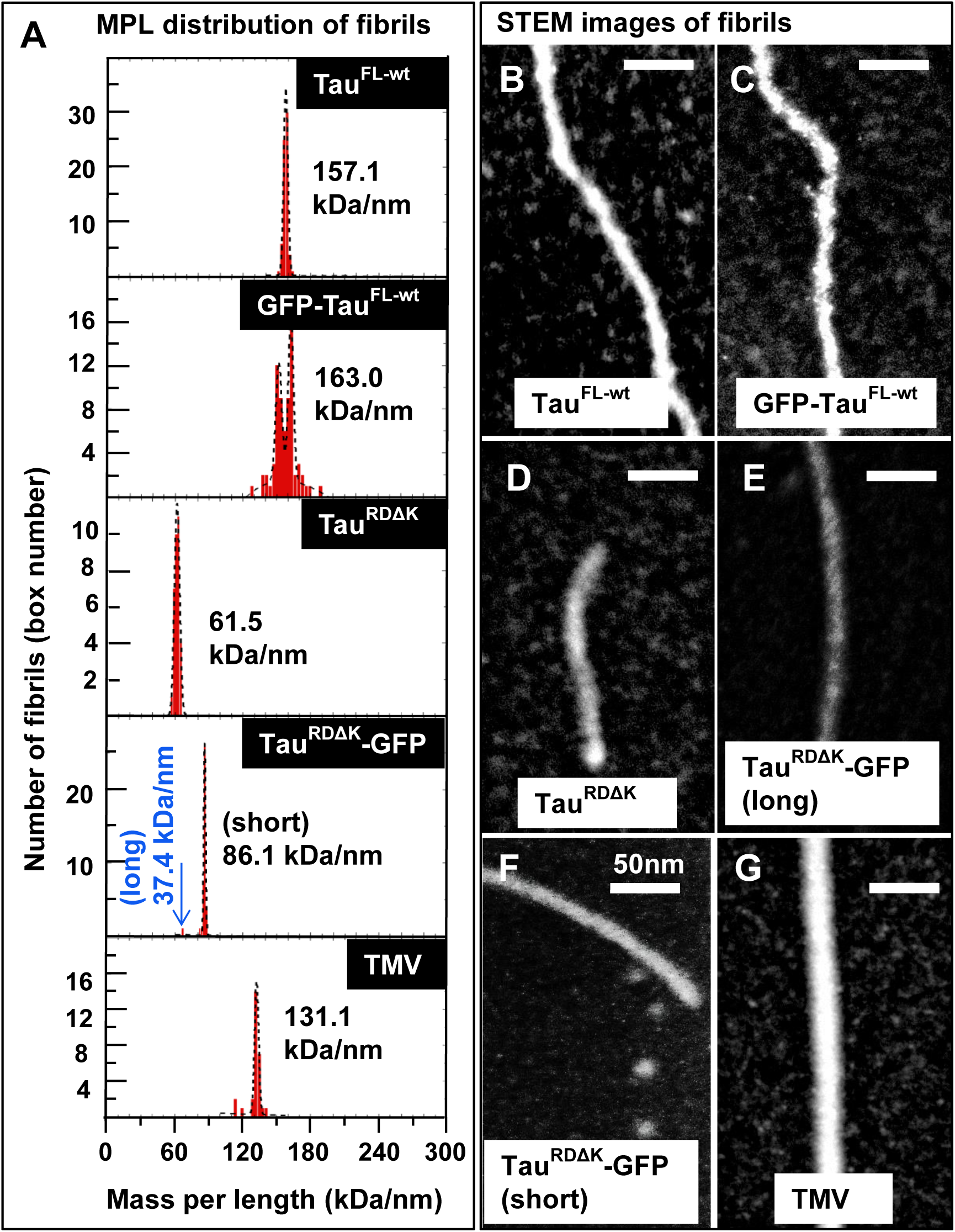
STEM imaging reveals that packing of Tau in assembly conditions is severely altered by GFP tags. **(A)** Mass per length histograms (MPL) of fibrils aggregated for 24 h at 37°C, with fitted peak values (in kDa/nm) listed for each protein. Examples of STEM images are shown for Tau^FL-wt^ **(B)**, GFP-Tau^FL-wt^ **(C)**, Tau^RDΔK^ **(D)**; Tau^RDΔK^-GFP fibrils (long) **(E)** and Tau^RDΔK^-GFP fibrils (short) **(F)**. MPL measurements were calibrated with tobacco mosaic virus (TMV, shown in the bottom histogram in **A** and in image **G**). The estimated molecules per nm are presented in table 1. Note that Tau^RDΔK^-GFP fibrils have been observed mostly as short and stubby fibrils, but that the long fibrils show a packing, less dense than the short fibrils.

## Discussion

The progression of Alzheimer Disease can be subdivided into several stages, based on abnormal changes in the neuronal protein Tau (notably hyperphosphorylation and aggregation) which spread in the brain with a predictable spatio-temporal sequence, following axonally connected pathways (Braak and Braak, 1991, Braak and Braak, 1996). This suggests that the signal of toxicity spreads via interconnected neurons and/or cells closely associated with them. Various modes of transmitting a toxic signal between cells can be envisaged (Brundin et al., 2008, Walsh and Selkoe, 2016). Currently one of the favored mechanisms is based on the “prion-like” spreading of Tau protein, whereby a misfolded and aggregation-prone form of Tau is transferred from a donor to an acceptor cell where it nucleates the conversion of native Tau to a misfolded state and thus causes aggregation (Vaquer-Alicea and Diamond, 2019, Holmes and Diamond, 2017). This is consistent with the fact that Tau is a neuron-specific protein, and that only neurons develop the abnormal changes of Tau that show up as neurofibrillary tangles or neuropil threads, consisting of bundles of paired helical filaments (PHF) or straight fibers of Tau (SF) (Fitzpatrick et al., 2017). A variant of the hypothesis is that the transfer of pathogenic Tau may occur not directly from donor to acceptor neuron, but indirectly via microglia and exosomes (Asai et al., 2015).

In the “prion-like” hypothesis, the concept of “seeding” plays an important role: In the strict sense it refers to the nucleated self-assembly of native tau in the acceptor cell onto the template provided by the incoming tau, thereby forming tau polymers with a misfolded conformation (Fig. 1). More generally it denotes the transfer of a pathogenic species (“misfolded” Tau) from donor to acceptor cell. The tacit assumption is that some species in this transformation is pathogenic for the acceptor cell (e.g. Tau polymers or oligomers). The main experimental procedure to test the pathogenic potential of a Tau preparation (e.g. from AD brain) is based on a reporter cell expressing the Tau repeat domain (∼13 kDa) fused to XFP [GFP or variants] (Holmes et al., 2014). Exposing the reporter cell to the extracellular Tau preparation may cause a local accumulation of fluorescence (or FRET, if Tau^RD^ is fused to FRET pairs like CFP/YFP). By analogy with the “prion-like” hypothesis, such inclusion with elevated fluorescence or FRET are interpreted as bona fide PHF-like aggregates, resulting from the internalization of the extracellular Tau and subsequent templated assembly of the Tau^RD^-XFP molecules.

We argue that this is an over-interpretation of the fluorescence data on structural grounds. As shown in this paper (Fig. 6, 7), steric hindrance prevents Tau^RD^-fusion proteins to assemble into PHF-like fibers. In a proper PHF the core has a tightly packed cross-β-structure, with a 0.47 nm distance between adjacent β-strands. However, this is not compatible with the size of an attached GFP molecule (size ∼2.4×4.2 nm, R_h_=2.8nm). Even if a templating Tau assembly were to reach the cytosol of an acceptor cell, it could not combine with the reporter Tau^RD^-GFP fusion molecules to propagate PHF-like structures. This situation is similar to other cases where an amyloidogenic protein fused to a compact reporter molecule lost its ability to aggregate into amyloid fibers (e.g. PABPN1-CspB, Aβ-GFP) (Ochiishi et al., 2016, Buttstedt et al., 2010).

**Figure 7:**
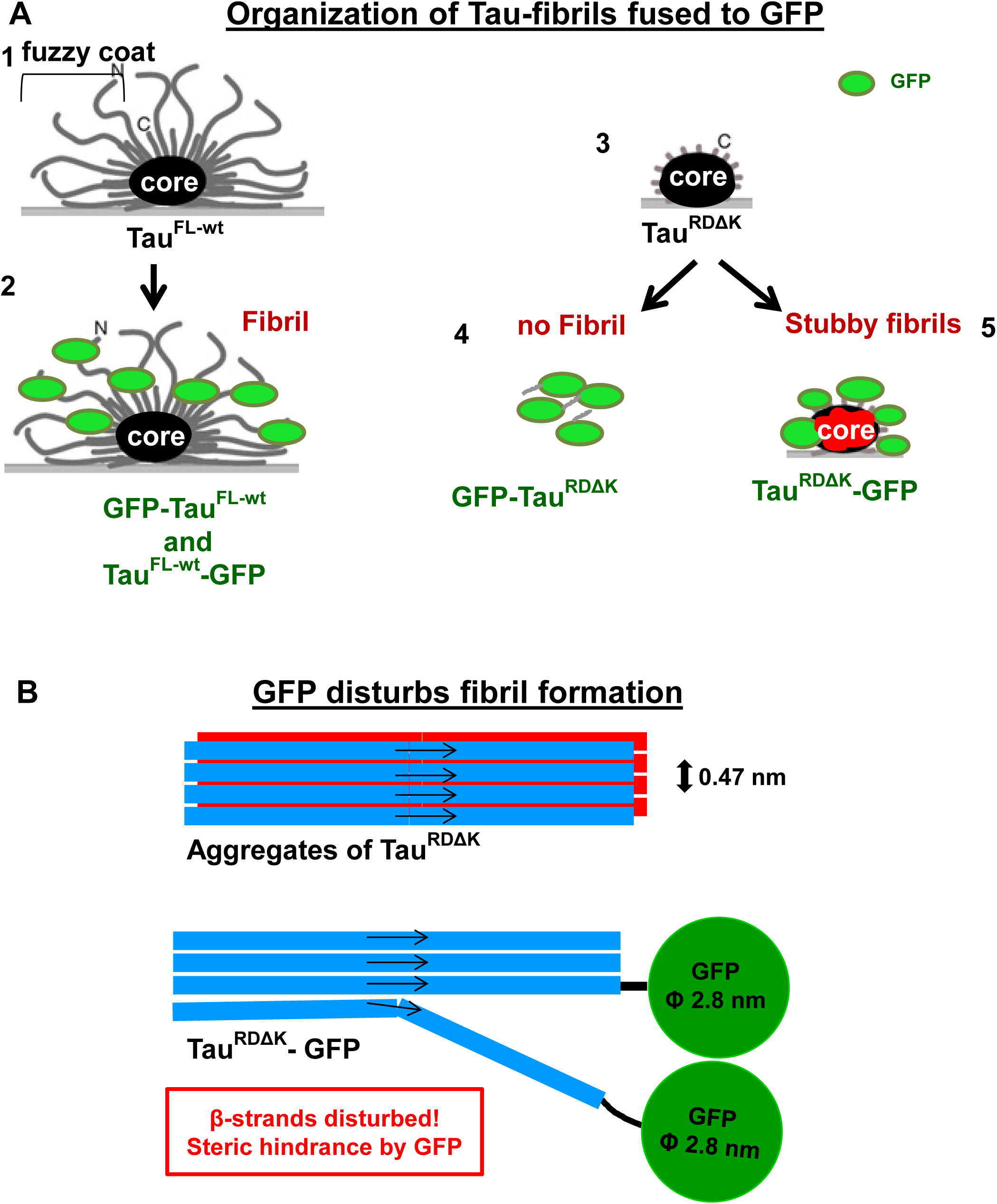
Models of organization of Tau fibrils fused to GFP. **(A)** Models of arrangement of GFP + Tau fusion constructs under aggregating conditions. The diagrams represent filaments lying on a support (grey line), viewed down the filament axis (black oval = core). (1, 2) Tau^FL-wt^ and GFP-Tau^FL-wt^ have a similar packing of their fibril cores (cross-β structure) with attached N- or C-terminal domains (1), plus GFP on either end (2). The PHF-like packing is possible because Tau is disordered and flexible, so that GFP molecules can be accommodated around the perimeter as a fuzzy “halo”. (3) Tau^RDΔK^ can form a PHF-like fibril core, but (4) GFP-Tau^RDΔK^ does not form any fibrils. (5)Tau^RDΔK^-GFP may assemble into short aberrant fibrils, but the arrangement is not compatible with that of AD-like filaments and results in perturbed core (red distorted structure). The GFP creates an outside layer (analogous to the fuzzy coat of PHFs), but the packing is less dense and the elongation is strongly disturbed. **(B)** Diagram of a side view of a PHF-like filament, made up of two parallel β-sheets (red and blue) with antiparallel orientations (arrows). The axial separation of β-strands is 0.47 nm. Size comparison of β-strand and attached GFP. If the GFP were attached sufficiently far away from repeat domain (as in FL tau), the amyloid core could be formed, with GFP molecules accommodated around the perimeter. If the GFP is too close to the β-strand core (as in Tau^RD^) this leads to steric hindrance which causes improper folding and disruption of the amyloid core.

How is it then possible to explain local accumulations of GFP-labeled Tau^RD^? One key factor is that the FRET effect extends up to distances of ∼10nm, considerably further than the spacing of secondary structures and sizes of the involved protein molecules, and it allows variable relative orientations of molecules. While full-length Tau tends to associate with microtubules, this is not the case for Tau^RD^ which has only a weak affinity for MT (Fig. 3) and instead is rather uniformly distributed in the cytoplasm. However, Tau^RD^ has a natively unfolded structure and can undergo multiple weak interactions with other hydrophilic or charged molecules. Examples are the interactions with ribosomes (Papasozomenos and Binder, 1987, Koren et al., 2019), stress granules (Apicco et al., 2018, Moschner et al., 2014), transport granules (Mercken et al., 1995, Konzack et al., 2007, Aronov et al., 2002). Moreover, several groups showed recently examples of how the low-complexity composition of Tau or Tau^RD^ enables it to become spontaneously enriched locally in phase-separated membrane-less compartments, particularly in combination with RNA or cytoskeletal proteins (Wegmann et al., 2018, Zhang et al., 2017, Hernandez-Vega et al., 2017, Ambadipudi et al., 2017).

The local inhomogeneities of Tau^RD^ distribution can be triggered by various extracellular stimuli independently of Tau, for example various stress signals, including oxidative stress or Aβ oligomers (Zempel et al., 2013). In particular, accumulations of GFP-labeled Tau^RD^ that mimic seeding can be triggered in neurons by cytokines such as TNFα released from activated microglia (Gorlovoy et al., 2009). It emphasizes the growing evidence that hallmarks of Tau pathology in neurons can be initiated by inflammatory signals from microglia (Ising et al., 2019). This can prompt both the aggregation of assembly-competent Tau as well as its hyperphosphorylation (e.g. by reducing the activity of phosphatases). Noteworthy in this context, “seeding” is observed primarily with heterogeneous Tau preparations that are tissue- or cell-derived, but is inefficient with purified tau samples.

How do these arguments affect approaches to therapy? Earlier attempts to prevent tau pathology were directed at pathological changes of Tau within neurons, assuming they were the carriers of toxicity. Examples are the inhibition of kinases or activation of phosphatases to reduce hyperphosphorylation (Medina, 2018), or inhibitor compounds to block aggregation (Wilcock et al., 2018, Wischik et al., 2014, Pickhardt et al., 2015, Pickhardt et al., 2007, Pickhardt et al., 2005) Overall these attempts were not successful. The “prion-like” hypothesis shifted the emphasis to the transfer of tau between cells, which could conceptually be intercepted by Tau-specific antibodies (Yanamandra et al., 2013, Congdon et al., 2016, Gu et al., 2013, Novak et al., 2017). Despite some encouraging results in transgenic mice, success of this approach is still uncertain and would not be expected if tau protein is not the carrier of pathogenicity, as suggested here.

## Abbreviations

AD: Alzheimer disease
AFM: atomic force microscopy
CNS: central nervous system
FRET: fluorescence resonance energy transfer
GFP: green fluorescent protein
PHF: paired helical filament
Tau^FL^: full-length Tau protein, largest isoform in human CNS (Uniprot P10636-F, htau40)
Tau^FLΔK^: full-length Tau protein with pro-aggregant mutation ΔK280
Tau^RD^: Tau repeat domain (amyloidogenic domain)
Tau^RDΔK^: Tau repeat domain with pro-aggregant mutation ΔK280
Sf9: cell line from Spodoptera frugiperda
STEM: scanning transmission electron microscopy
EM: electron microscopy
MPL: mass per length
Wt: wildtype

## Acknowledgements

We thank Ms. Carola Troger for her assistance with STEM experiments and the CAESAR electron microscope facility. This project was supported by funding from DZNE and MPG.

## Supplementary figure legends

**Table. S1: Amino acid sequences of tau constructs**.

Amino acid sequences of the tau constructs used in this study. See also figure 2 (A-B). A linker sequence was introduced to separate the GFP from the tau proteins sequence in the repeat domain Tau constructs. Sequences in green color are GFP protein sequences; blue are linker sequences and red are his tag sequences.

**Fig S1:**
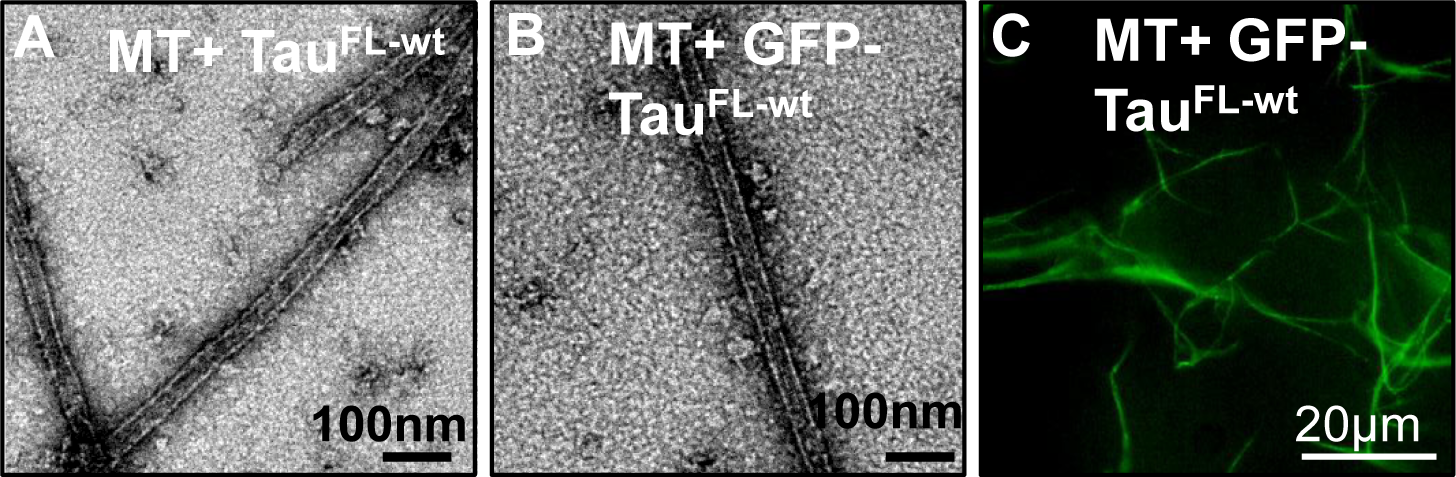
Full length GFP-Tau^FL^ promotes assembly of microtubules in vitro. **(A)** Microtubules (10 µM) assembled in vitro in presence of untagged Tau^FL-wt^ (5 µM) or **(B)** GFP-Tau^FL-wt^ (10 µM) were imaged by negative stain electron microscopy. Both Tau constructs efficiently promote the assembly of typical microtubules. **(C)** Fluorescence microscopy image of microtubules assembled in vitro with GFP-Tau^FL-wt^ reveal the distribution of the GFP tag along the length of microtubules.

## Materials and Methods

### Materials

– All chemicals were obtained from Sigma-Aldrich (Taufkirchen, Germany), Fluka (Seelze, Germany), and Roth (Karlsruhe, Germany) in highest purity if not stated otherwise. Heparin 5000 or 16000 was purchased from Fisher Scientific (#BP2524-10). MSNL-10 probe for AFM measurements were from Bruker, USA.

### Cells and viruses

Sf9 cells were obtained from Invitrogen (Life Technologies, DE) and grown at 27°C in monolayer culture with Grace’s medium (Sigma /Life Technologies TM) supplemented with 10% fetal bovine serum and 1% penicillin / streptomycin (PS).

Sapphire Baculovirus DNA was obtained from Orbigen/Biozol (Eching, DE) and pVL1392 from Invitrogen (Life Technologies, EM). MultiBacTurbo system contains the acceptor plasmid pACEBac1 and the E.coli strain DH10MultiBac /Turbo contains the acceptor bacmid. MultiBacTurbo baculoviral DNA were obtained from EMBL Grenoble Outstation(Bieniossek et al., 2012).

### Plasmids and baculovirus construction

The hTau40 cDNA, the longest Tau isoform in human CNS (2N4R), was tagged with 6xHis at the C-terminus using PCR amplification and the modified Tau-His cassette was introduced into pET3a plasmid (Novagen, DE) linearized with NdeI enzyme succeeding in pET3a/htau40/6xHis plasmid for the protein expression in E.coli. The cDNA cassette of GFP-hTau40 (consisting of M1 – L239 of GFP sequence bridged by one aa: His to htau40 (M1 – L441) sequence elongated with 6x His at the C-terminus was inserted into pET3a plasmid succeeding in E.coli expression vector pET3a GFP/htau40/6xHis.

The cDNA cassette of htau40-GFP (consisting of htau40 (M1-L441) bridged by 13 aa AH linker sequence: GAPGSAGSAAGSG to M1-L239 GFP terminated with 6xHis Tag was also inserted into pET3a plasmid leading to the pET3a htau40/GFP/6xHis expression vector for E.coli. Similarly, the cDNA cassette of GFP-Tau^RDΔK^-6xHis consisting of GFP, pro-aggregant Tau repeat domain (4-repeat construct K18, M-Q244-E372, with the deletion mutation ΔK280 terminated with 6xHis Tag (GFP-Tau^RDΔK^) was also inserted into pET3a plasmid, yielding the pET3a GFP-Tau^RDΔK^-6xHis expression vector. This mutation strongly accelerates aggregation due to pronounced β propensity (von Bergen, et al., 2001).

A second group of proteins: Tau^RDΔK^ –GFP and GFP/6xHis proteins was expressed in the baculovirus expression system. A cDNA cassette containing Tau^RDΔK^ tagged with GFP at its C-terminus (Tau^RDΔK^–GFP) followed by a 6xHis tail was inserted into pVL1392 vector resulting in the plasmid pVL1392 Tau^RDΔK^/GFP/6xHis. After mixing with Saphire TM baculovirus DNA, this plasmid was used for the generation of baculovirus and protein expression in Sf9 insect cells as described before (Biernat et al., 1993).

The GFP cDNA sequence fused to a 6xHis tail at its C-terminus was inserted into the pACEBac-1 acceptor vector to generate the pACEBac/GFP/6xHis plasmid. This plasmid was subsequently subjected for the Tn7-dependent integration into the baculoviral genome of DH10 MultiBacTurbo E.coli cells for the generation of baculovirus encoding GFP-His-tagged protein.

The purified MultiBacTurbo bacmid encoding GFP protein was transfected into Sf9 cells for the generation of baculoviruses and subsequent protein expression (Bieniossek et al., 2012). An overview of the Tau constructs is shown in Fig. 2 (for sequence see Supplement Table S1).

### Protein preparation and purification

Proteins were expressed either in E.coli or in insect Sf9-cells using the baculovirus expression system. The Tau proteins hTau40wt (2N4R) and Tau^RDΔK^ (K18 construct, (M) Q244-E372 with deletion of K280) were expressed in E.coli and purified as described earlier (Barghorn et al., 2005).

The proteins Tau-His (hTau40/6xHis; 2N4R), GFP-Tau^RDΔK^ (GFP- (M)Q244-E372 with deleted K280/6xHis) and GFP-Tau (GFP-hTau40/6xHis; 2N4R) and Tau-GFP (hTau40/-GFP/6xHis; 2N4R) were expressed as fusion proteins with 6x polyHis tail at the C-terminus in E.coli strain BL21(DE3) (Merck-Novagen, Darmstadt). After harvesting, the E.coli bacteria cell pellet was directly re-suspended in lysis buffer [50 mM Tris HCl pH 7.3, 300 mM NaCl, 10% glycerol, 0.5 mM TCEP, 10 mM Imidazol, 1 mM Benzamidin, 1 mM PMSF and 10µg/ml each of protease inhibitors leupeptin, aprotinin, and pepstatin], in the ratio 1g E.coli pellet to 10 ml lysis buffer and disrupted by French press.

Protein Tau^RDΔK^-GFP (M) Q244-E372 with deletion of K280-GFP/6xHis), GFP/6xHis and also GFP-Tau (GFP-hTau40/6xHis; 2N4R) and (hTau40/-GFP/6xHis; 2N4R) were expressed in Sf9 insect cells from Invitrogen. Sf9 cells were infected with recombinant virus at a MOI of > 1 typically in six T150 cell culture flasks containing 75% confluent Sf9 cells. Cells were incubated for three days at 27°C, collected and re-suspended for preparation in the lysis buffer [50 mM Tris HCl pH 7.3, 300 mM NaCl, 10% glycerol, 0.5 mM TCEP, 10 mM Imidazol, 1 mM Benzamidin, 1 mM PMSF and 10 µg/ml each of protease inhibitors leupeptin, aprotinin, and pepstatin], in the ratio 1g Sf9 pellet to 10 ml lysis buffer and disrupted by French press.

The lysates of the E.coli bacteria and Sf9 cells were cleared by centrifugation in a Beckman Optima L-80 XP Ultracentrifuge with a Ti45 rotor (40,000 rpm. for 60 min at 4 °C), applied to Ni2+ ion affinity chromatography His Trap FF column (GE Healthcare), and purified using an Äkta pure chromatography system (GE Healthcare). Following extensive wash with 12 column volumes (CV) of wash buffer (50mM Na phosphate buffer pH 7.2, 300 mM NaCl, and 25 mM imidazole) the protein was eluted with elution buffer (50mM Na Phosphate buffer pH 7.2, 300 mM and 1 M imidazol).

If necessary the protein breakdown products were separated in the second chromatography step using gel filtration on a Superdex G200 column (GE Healthcare, Freiburg). PBS buffer was used for gel filtration column (137 mM NaCl, 3 mM KCl, 10 mM Na2HPO4, 2 mM KH2PO4, pH 7,4) with 1 mM DTT (freshly added).

### Polymerization of Tau

#### Light scattering

Tau aggregation was monitored by 90° angle light scattering at 350nm in a FluoroMax spectrophotometer (HORIBA). 50 µM Tau protein in suspended in BES buffer, pH 7.0 (20 mM BES, 25 mM NaCl) and supplemented with 12.5µM heparin 16000 or heparin 5000. Heparin 5000 and 16000 induce the aggregates at the same level and there is no difference between the aggregates formed under these conditions. This mixture was incubated at 37°C. At different time points 20µl of the samples were analyzed by light scattering at 350 nm and then the samples were brought back to original tube for further aggregation.

### Sedimentation assay and western blot

The aggregated samples (t=48 h) were sedimented at 61,000 rpm (TLA 100.3 rotor) for 1h at 4°C. The supernatant was collected and the pellet was resuspended to the same volume as supernatant. 5µl of the samples were resolved on 10% SDS gel and immunoblotted as described in (Kaniyappan et al., 2017). Later the blot was probed with K9JA (1:5000 dilution) and GFP (1:1000 dilution) antibodies and the signal was detected by chemiluminescence method.

### Turbidity assays for MT assembly

Tau-induced microtubule assembly was monitored by 90° angle light scattering at 350nm in a FluoroMax spectrophotometer (HORIBA). 10 µM PC-purified Tubulin were mixed with 5 µM Tau protein in RB-Buffer (100 mM PIPES pH 6.9, 1 mM DTT, 1mM MgSO4, 1mM EGTA, 1 mM GTP). The polymerization was started by transferring the ice-cold Tubulin-Tau-solution to the 37°C warm cuvette-holder and the reaction started once the temperature was reached. Tubulin assembly was monitored for 15 min.

### Mass per length analysis by STEM

#### Sample preparation

Holey carbon grids (Quantifoil R2/1) covered with either 2 nm amorphous carbon (Quantifoil, R2/1+2 nm C) or graphene were used for sample preparation. The preparation of graphene grids was done according to a method described earlier (Pantelic et al., 2011).The sample concentration was adjusted for MPL measurements to a final concentration of 5–10 molecules / grid hole. The protein solution was mixed with Tobacco mosaic virus (TMV) prior to application to the grids. The TMV was used to calibrate the mass measurements. 10 µl sample was applied to the grids for 2 min. Excess liquid was blotted away using filter paper. Samples were washed 3 times with doubled distilled water to remove buffer salts and afterwards air dried or frozen and vacuum dried at – 80 °C for 12 h.

### Scanning-transmission electron microscopy (STEM)

For the MPL experiments, a Zeiss Libra200 MC Cs-STEM CRISP (Corrected Illumination Scanning Probe) was used. The instrument was operated at 200 kV. The CRISP is equipped with a monochromated Schottky-type field emission cathode and a double hexapole-design corrector for spherical aberrations of the illumination system (Cs-corrector). A high-angle annular dark field (HAADF) detector (Fischione Instruments, USA) was used for imaging. The images were recorded at a convergence angle of 16 mrad and an acceptance angle of 20 mrad. Images were recorded at a pixel size of 0.6 nm with a dwell time of 70 µs/px. The total dose per image was 400 e/nm2. Mass determination was done using the software PCMASS32 (Schutz et al., 2015). In a first step images were calibrated using tobacco mosaic virus as calibration sample. Filament regions were manually selected and masked. The full datasets were plotted as histograms and fitted to Gaussian curves.

### Transmission electron microscopy (TEM)

Samples for negative stain transmission electron microscopy were placed on 200 mesh formvar-carbonated copper grids with (Plano, Quantifoil #S162). Grids were glow-discharged for 30 s (Baltec, MED010) prior to sample addition. The protein solution was removed after 3 min incubation time by filter-paper and the grids washed 3 times with ddH2O (grid on top of each drop) to remove buffer salts, then followed by staining with 2% uranyl acetate for 60 seconds and a finally removing the stain slowly with a wet-filter paper and rapid air drying. To prepare grids with microtubules for TEM, all steps were carried at out at 37°C and with pre-warmed solutions. All specimens were analyzed at 200kV using a JEOL JEM-2200FS TEM.

### Atomic force microscopy (AFM)

For AFM measurements, Tau fibril samples were diluted in PBS buffer for a final concentration of 0.5 – 1µM. 30µl of the sample was placed onto freshly cleaved mica and allowed to adsorb for 10 minutes at room temperature. Unattached and excess proteins were removed by rinsing the sample with PBS for 4-5 times. Finally, the sample on the mica was filled with imaging buffer (10 mM Tris-HCl, pH 7.4, 50 mM KCl). AFM imaging was performed in oscillation mode using JPK NanoWizard® ULTRA Speed AFM system and MSNL-10 probe with “F” cantilever. The amplitude set point and the gains were adjusted manually to control the thermal drift and to achieve the minimal force between the cantilever and the sample. AFM images were processed by JPK data processing software. Fibril heights/widths were measured by the inbuilt cross-sections method of JPK data processing software. For the analysis of fibril width 50 fibrils per condition were measured.

---

**Table S1.**
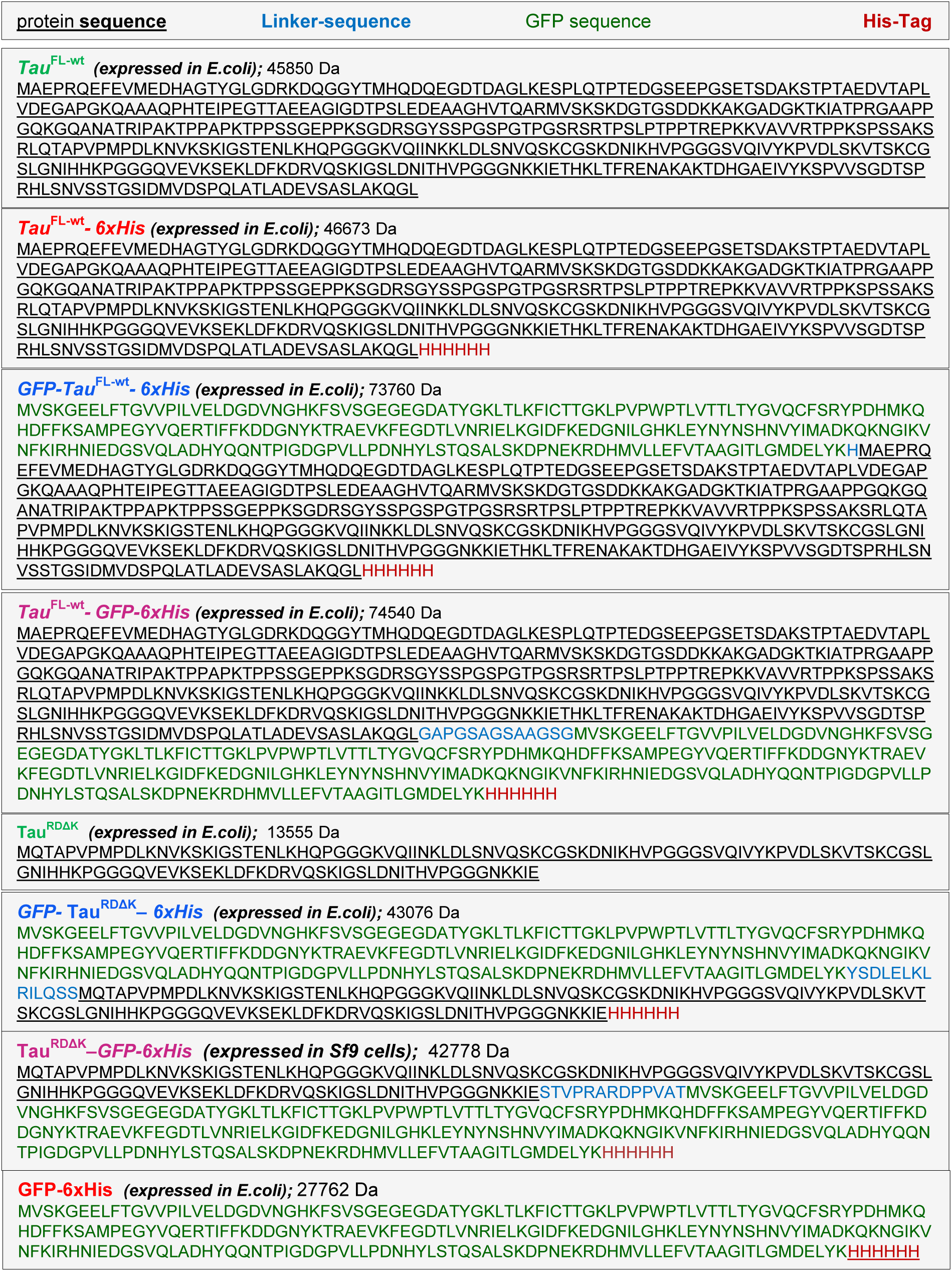
Amino acid sequences of the tau constructs used in this study. See also Figure 1 (A-B). A linker sequence was introduced to separate the GFP from the tau proteins sequence in the repeat domain Tau constructs.

